# Does fatigue influence joint-specific work and ground force production during the first steps of maximal accelerative running?

**DOI:** 10.1101/2022.04.21.489102

**Authors:** Shayne Vial, Jodie Cochrane Wilkie, Mitchell Turner, Mark Scanlan, Anthony J. Blazevich

## Abstract

The rate of initial acceleration during the first steps of a maximal-effort (sprint) run often determines success or failure in prey capture and predator evasion, and is a vital factor of success in many modern sports. However, accelerative events are commonly performed after having already run considerable distances, and the associated fatigue should impair muscle force production and thus reduce acceleration rate. Despite this, the effects of running-induced fatigue on our ability to accelerate as well as the running technique used to achieve it has been incompletely studied. We recorded 3-D kinematics and ground reaction forces during the first three steps of the acceleration phase from a standing start before and after performing a high-speed, multi-directional, fatiguing run-walk protocol in well-trained running athletes who were habituated to accelerative sprinting. We found that the athletes were able to maintain their rate of initial acceleration despite changing running technique, which was associated with use of a more upright posture, longer ground contact time, increased vertical ground reaction impulse, decreased hip flexion and extension velocities, and a shift in peak joint moments, power, and positive work from the hip to the knee joint; no changes were detected in ankle joint function. Thus, a compensatory increase in knee joint function alleviated the reduction in hip flexor-extensor capacity. These acute adaptations may indicate that the hip extensors (gluteal and hamstring muscle groups) were more susceptible to fatigue than the ankle and knee musculature, and may thus be a primary target for interventions promoting fatigue resistance.

## Introduction

The top running speed of a terrestrial animal is commonly considered to influence its capacity to hunt prey or avoid capture, and is often successfully used by animals such as cheetah, antelope, wildebeest, and others (Wilson et al., 2018). However, even the fastest animals frequently fail in their attempt to capture prey (Wilson et al., 2020), and top speeds may never be reached in a confrontation by either predator or prey, even by the fastest animals (Brown et al., 1995; Howland, 1974; Humphries and Driver, 1970; Wilson et al., 2013). Alternatively, the ability to accelerate, including the ability to abruptly change direction, typically dictates a successful outcome more so than the top speed (Husak and Fox, 2006; Irschick et al., 2005; Wheatley et al., 2018; Wilson et al., 2013). Indeed, the early identification of a threatening stimulus and, importantly, the initial movement reaction speed, is considered to be key to overall success in situations where acceleration rate strongly influences the chase outcome (Isbell, 1994; Isbell et al., 2018; Quigley and Herrero, 2009). Thus, the acceleration capacity (linear or non-linear) of an animal, particularly within the first steps, could be more critical to survival than top running speed. In modern sports, a similar observation is made, with many athletes successfully capturing or evading an opponent through rapid acceleration, regardless of whether changes of direction occur, without reaching maximum running speed (Gabbett, 2010; Varley et al., 2012). Therefore, the capacity for skeletal muscles in both humans and other animals to produce the substantial forces required to accelerate the body during the early phase of sprinting will critically influence outcome success.

Modern human athletes frequently shift between walking, jogging, and sprinting gaits during a competition or match (Brewer et al., 2010; Gabbett, 2010; Johnston et al., 2012). As the competition progresses, e.g. in the latter stages of a quarter or half in team sports, fatigue may become prominent (Ekstrand, 2008; Woods et al., 2004). Yet, the demands of the game remain similar and athletes are still required to accelerate rapidly to catch or evade an opponent, requiring substantial physical exertion whilst fatigued (Gabbett, 2004; Harper et al., 2019; Malone et al., 2018; Mohr et al., 2004; Varley and Aughey, 2013). Early human hunter-gatherers too would have covered vast distances by walking and jogging before sometimes needing to accelerate rapidly to capture prey or evade predation (Liebenberg, 2006; Liebenberg, 2008; Quigley and Herrero, 2009). As fatigue deepens, muscle force generation capacity becomes impaired and our maximum acceleration rate consequently decreases (Ciacci et al., 2017; Hautier et al., 2000; Nagahara et al., 2014; Pinniger et al., 2000; Small et al., 2009), increasing the risk of failure to either catch an opponent or prey or to evade capture by another, which would be costly to both modern athletes and our human ancestors alike.

The physical proportions of humans, with relatively shorter arms, longer legs, and larger hip-extending gluteal muscles compared to our earlier ancestors, may have played a role in allowing humans to shift between movement patterns such as rapidly accelerating to reach high running speeds, changing the direction of running (e.g. 45° turn from direction running), jumping, and climbing over obstacles (Liebenberg, 2006; Pickering and Bunn, 2007; Steudel-Numbers and Wall-Scheffler, 2009). The ground reaction forces produced result from the joint forces and moments generated by muscles surrounding lower limb joints. But because each muscle has a different function, morphology, location, and thus contraction force and/or velocity capacity (Barry and Enoka, 2007; Pinniger et al., 2000), muscles can fatigue to different magnitudes and at different rates (Hautier et al., 2000; Lord et al., 2018). Thus, fatigue-related torque degeneration may be muscle- or joint-dependent (Dutto and Smith, 2002; Jones, 2010; Lord et al., 2018). Therefore, to retain the same movement velocity as before fatigue, muscle force (work) and power production may be shifted from highly fatigued muscles to other, less fatigued muscles, which may then act across different joints. This would confer a change in movement pattern and joint-specific contribution to the task, although such changes may mitigate acceleration loss and thus provide the greatest chance of movement success, i.e., rate of acceleration (Gates and Dingwell, 2011; Hiemstra et al., 2001). While the mechanism of muscle force redistribution during fatigue in rapid, accelerative running is plausible, it remains untested. Furthermore, it is not known whether any changes in relative joint contributions to total work would be detrimental to acceleration performance or increase the risk of injury due to altered joint and soft tissue loading. A better understanding of fatigue-induced technique alterations may allow for more specific hypotheses to be drawn and thus for future research to be designed. Currently, we know relatively little about how humans move in the initial few steps of maximal accelerative running, or how our movement strategies are influenced by fatigue.

Given the above, the primary purpose of the current study was to describe the kinematic and kinetic patterns of the dominant (stronger) and non-dominant (weaker) legs in the first three steps of both non-fatigued and fatigued accelerated running (sprinting), and thus to assess the changes in within-limb distribution in response to fatiguing running exercise. The dominant and non-dominant legs were not directly compared because the velocity of the centre of mass (CoM) was different between steps and a true comparison was not possible. Therefore, we tested the hypothesis that reduced horizontal velocity and substantial, acute kinematic and kinetic adaptations would be observed with fatigue.

## Methods

### Population and training history

Thirteen semi-professional male Association Football (Soccer) players (age: 19.1 ± 2.1 y, body mass: 72.5 ± 6.9 kg, height: 175.0 ± 7.7 cm) participated in this study. This cohort was chosen because they undertook regular training and participated in matches that include walking, slow jogging, and running at varying speeds. To ensure use of their own freely chosen running technique, inclusion criteria included the participants performing no previous formal running technique instruction or training. No subjects participated in formal strength training or other supplementary training (e.g., plyometrics, etc.) that might influence their running performance, technique, or response to fatiguing running exercise. All participants were free from injury for at least six months before testing, reported no residual impediments from previous injury, and wore their normal training attire, which included the same running shoes during testing. Only outfield players were accepted into the study as goalkeepers tend not to (or rarely) perform maximal sprinting during training or match play. The study was approved by Edith Cowan University of Human Ethics Committee and players gave written consent prior to testing.

### Biomechanical measurements

Thirteen VICON motion capture cameras (Oxford Metrics Ltd., Oxford, UK) set at a frame rate of 250 Hz were positioned to capture the region from 0 to 5 m of the linear running acceleration trials. Ground reaction force data were simultaneously collected by five in-ground 600 × 900-mm triaxial force platforms positioned in a series (Kistler Quattro, Type9290AD, Victoria, Australia) sampling at 1000 Hz. Motion capture cameras were arranged around the in-ground force platforms over which the subjects performed their running acceleration trials. The subjects started behind the first force platform and the finish line was located 20 m from the start to ensure that subjects did not decelerate through the data capture zone; that is, although only the first three steps were examined, the subjects were required to accelerate maximally over a longer distance, which might more likely lead to opponent capture or evasion.

### Protocol

Upon arrival, height and body mass were recorded for each subject, and a custom cluster-based set of retroreflective markers were placed on anatomical landmarks for 3-dimensional motion capture (see Supplementary File for details). Subjects then completed a standardised, comprehensive warm-up (also described in Supplementary File) which included practice trials for single-leg vertical jumps (SLVJ) on each leg and acceleration run efforts performed at 50%, 75%, and 100% of maximal effort for the full 20-m distance from a standing stationary start; the subjects started in a semi-crouched position with one foot in front of the other and the toes of the front foot on the start line; the hands did not touch the ground in this position.

Once the warm-up and practice trials were completed, three SLVJs were performed on each leg to obtain jump heights to determine dominant and non-dominant legs. The SLVJ test was used to define the ‘strongest’ (dominant) and ‘weakest’ (non-dominant) limbs as it encapsulates the ability to rapidly produce force without influence from the other leg. Therefore, the leg that produced greater jump height was selected as dominant. In some studies, researchers have designated the preferred kicking leg as dominant, which may not be the stronger of the two legs and thus may not accord with our definition. Then, once in their standing start position, a countdown from 5 to 1 led into the first 20-m maximal acceleration trial (see Figure 1 below), and subjects repeated this twice to obtain the *pre-fatigue* measurements. Verbal encouragement was given to ensure the subjects accelerated maximally and did not decelerate through the data capture zone. Subjects were allowed exactly 60 s of rest between trials, which included walking back to the start line. This relatively short rest was used to minimise the recovery from fatigue in post-running (fatigued) trials but did not induce detectable running fatigue in the non-fatigued tests (see Results).

**Figure 1.**
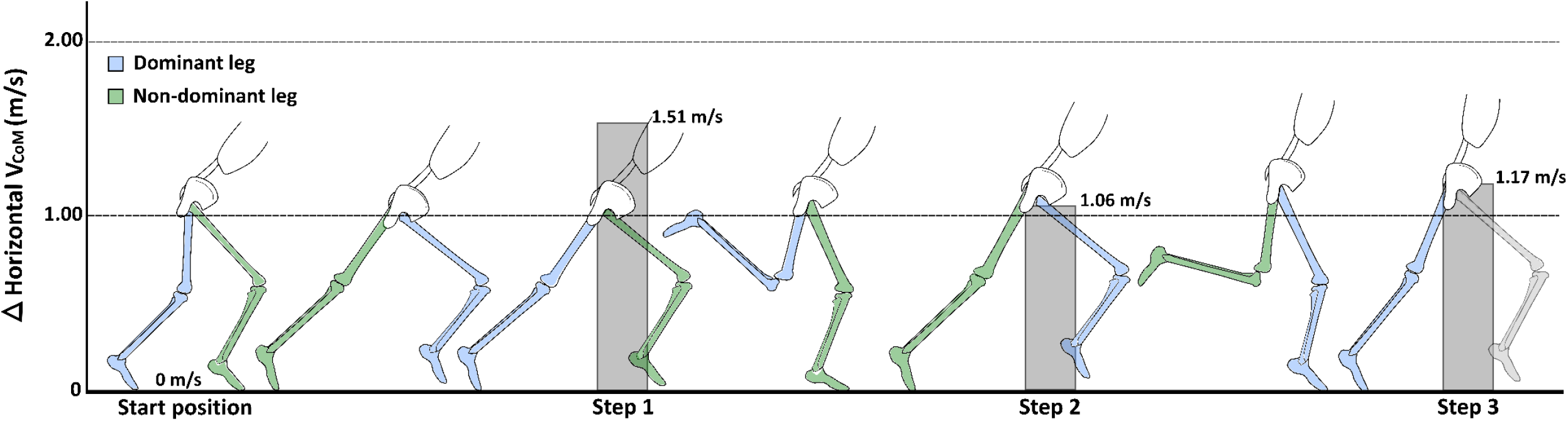
The first three steps of accelerated running from a standing start overlayed on the step-to-step change in horizontal centre of mass velocity (V_CoM_). The vertical axis shows the change in V_CoM_ from Step 1 to Step2 and Step 3. The greatest change in horizontal V_CoM_ was observed in Step 1, followed by Step 3, with Step 2 being the lowest. (Blue) DL: dominant leg; (Green) NDL: non-dominant leg. The step cycle was defined from the second half of the swing phase to toe-off. Each step was defined from the second half of the recovery phase through to toe-off

Once the first set of maximal acceleration trials was completed, the subjects jogged for 100 m at a comfortable pace to a grassed sports surface where markers placed on the surface demarcated a soccer-specific fatiguing exercise protocol (Ball–Sport Endurance and Sprint Test; BEAST 45) that would last 45 min (Williams, Abt, & Kilding, 2010; see Appendix). The fatigue protocol has been shown to be a valid and reliable simulator of soccer match play with respect to the movement patterns (walking, running, jumping, backwards running), intensity (i.e. distance and speed) and fatigue levels observed during a match (see Supplementary File) (Delextrat et al., 2018; Matthews et al., 2017; Williams et al., 2014; Williams et al., 2010). This protocol was familiar to the subjects, who were practiced in completing it without a notable pacing strategy and did not require further, extensive familiarisation. The protocol requires all directions of movement that might be needed in the chase or evasion of an agile animal and is also reflective of the movement patterns required in many modern sports. Subjects were also verbally instructed to run as fast as possible during the sprint sections and to decelerate within the allocated areas. After performing the fatiguing exercise, subjects jogged at a comfortable pace (100 m) back to the laboratory, performed three SLVJs on each leg, and then returned to the start line before a countdown from 5 to 1 led into the first of three maximal acceleration trials (post-fatigue test; *fatigued* trials) with 60 s of walk-back recovery between trials.

### Data analysis

All static and dynamic motion capture trials were digitised using VICON Nexus software (Oxford Metrics Ltd., Oxford, UK) and exported as C3D files to Visual 3D (C-Motion, Germantown, MD, USA). Ground reaction force and marker trajectory data were filtered using a fourth-order (zero lag), low-pass Butterworth filter with a 15-Hz cut-off frequency, as determined through residual analysis of marker trajectory data (van den Bogert & de Koning 1996). Together, subject height, body mass and the static standing trial data were used to create a skeletal model that included the trunk, pelvis, thigh, shank, and foot segments using Visual 3D software. Dynamic calibration trials were collected to determine functional joint centres for the hip, knee, and ankle joints. Data of all three non-fatigued and fatigued acceleration trials were averaged for each subject for the dominant and non-dominant limbs from the retraction and protraction phases for the first, second, and third steps of the acceleration phase, respectively. Each step was defined from the second half of the recovery phase through to toe-off (see Figure 1).

### Statistical Analysis

Paired t-tests were used to compare discrete kinematic, kinetic, and temporal variables between non-fatigued and fatigued conditions. All statistical analyses for discrete variables were performed using JAMOVI (Version 1.6, Sydney, Australia). For continuous data, Statistical Parametric Mapping (SPM; SPM1D open-source package, spm1d.org) in Python was used to compare non-fatigued and fatigued joint kinematics and kinetics for the first and third steps of the acceleration phase for the dominant leg (Pataky, 2012). These data were normalised to stride time, representing 0%-100% of the retraction and protraction phases for the first and third steps, respectively. Paired t-tests (SPM) were then completed separately at each of the 101 time points, providing a statistical parametric map (SPM{t}). If the SPM{t} surpasses the critical threshold with a successive collection of ≥5 points, a statistically significant difference was considered (Colyer et al., 2018) between non-fatigued and fatigued conditions. In all tests, the alpha level was set at 0.05.

## Results

The initial three steps of accelerated sprint running were compared between non-fatigued and fatigued conditions. The non-fatigued and fatigued within-limb comparisons included the dominant leg (DL; stronger) during the first and third steps and non-dominant leg (NDL; weaker) during the second step (see Figure 1).

### Step-to-step comparisons: Non-fatigued, accelerated running

The average horizontal centre of mass (CoM) velocity increased by ∼41% from Step 1 (S1) to Step 2 (S2) (1.51 m/s to 2.57 m/s) and then ∼31% from S2 to Step 3 (S3) (2.57 m/s to 3.74 m/s), while vertical CoM velocity and displacement (i.e., from maximum to minimum heights) remained constant. Ground contact times decreased by ∼9% from S1 to S2 and ∼5% from S2 to S3 (Table 1). In S1, the foot landed 0.05 m behind the body’s CoM, while foot contacts in S2 and S3 were both 0.09 m in front of the CoM (Figure 2). Horizontal foot velocity relative to CoM at ground contact increased with each successive step, i.e., increased forward foot speed relative to CoM, while vertical velocity, i.e., downward foot speed, did not change. During the steps, net horizontal impulse decreased by ∼22%, while net vertical impulse remained relatively constant (Table 1).

**Table 1.** Ground contact times across three steps during running acceleration trials before (non-fatigued) and after (Fatigued) fatiguing running. *Statistical difference (*t*-test) between non-fatigued and fatigued trials (p < 0.05).

|  | **Non-fatigued** | **Fatigued** | **Mean diff** | **95% CI (change)** |
| --- | --- | --- | --- | --- |
| Ground contact time (s) | **Mean ± SD** | **Mean ± SD** |  |  |
| Step 1 | 0.202 ± 0.02 | 0.201 ± 0.01 | -0.01 | 0.00, -0.01 |
| Step 2 | 0.183 ± 0.02* | 0.192 ± 0.01* | -0.01 | -0.01, -0.01 |
| Step 3 | 0.174 ± 0.01* | 0.184 ± 0.01* | -0.01 | -0.01, -0.01 |

**Figure 2.**
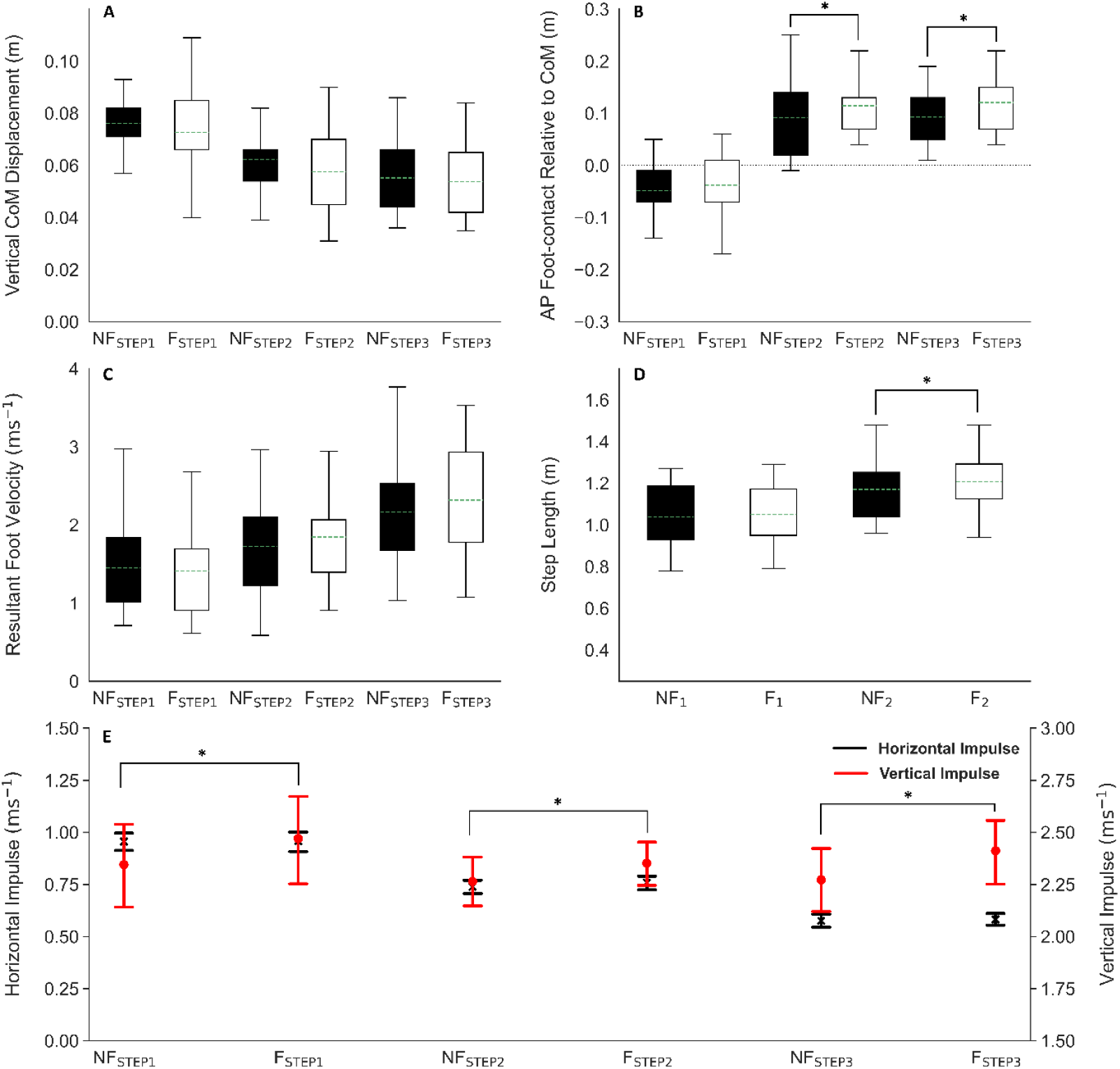
Non-fatigued and fatigued accelerated sprint running performance outcomes. After fatiguing exercise, the foot contacted the ground further in front of the CoM ((B) S2; S3), step length increased ((D) S2 only) and vertical impulse increased across all steps (E). NF, non-fatigued (black); F, fatigued (white); step number indicated in subscript. A) vertical CoM displacement (metres); B) anterior-posterior (AP) foot-ground contact distance relative to the horizontal position of CoM (metres); C) resultant foot velocity (horizontal and vertical; metres per second); D) step length (metres); E) relative net horizontal impulse (ms^-1^; primary axis) and relative net vertical impulse (ms^-1^; secondary axis). *Statistical difference (*t*-test) between non-fatigued and fatigued trials (p < 0.05).

### Step-to-step kinematic, kinetic, and mechanical work comparisons: Non-fatigued, accelerated running

The sagittal trunk angle relative to the ground at ground contact increased in each successive step (∼58° for S1, ∼72° for S2, and ∼84° for S3). On average, peak hip flexion angle tended to decrease slightly with each successive step (see Figure 3), while peak knee extension angle remained constant for S1 and S3 but decreased slightly (greater flexion) between S1 and S2 (see Figure 4). No differences were observed in ankle joint kinematics between steps.

**Figure 3.**
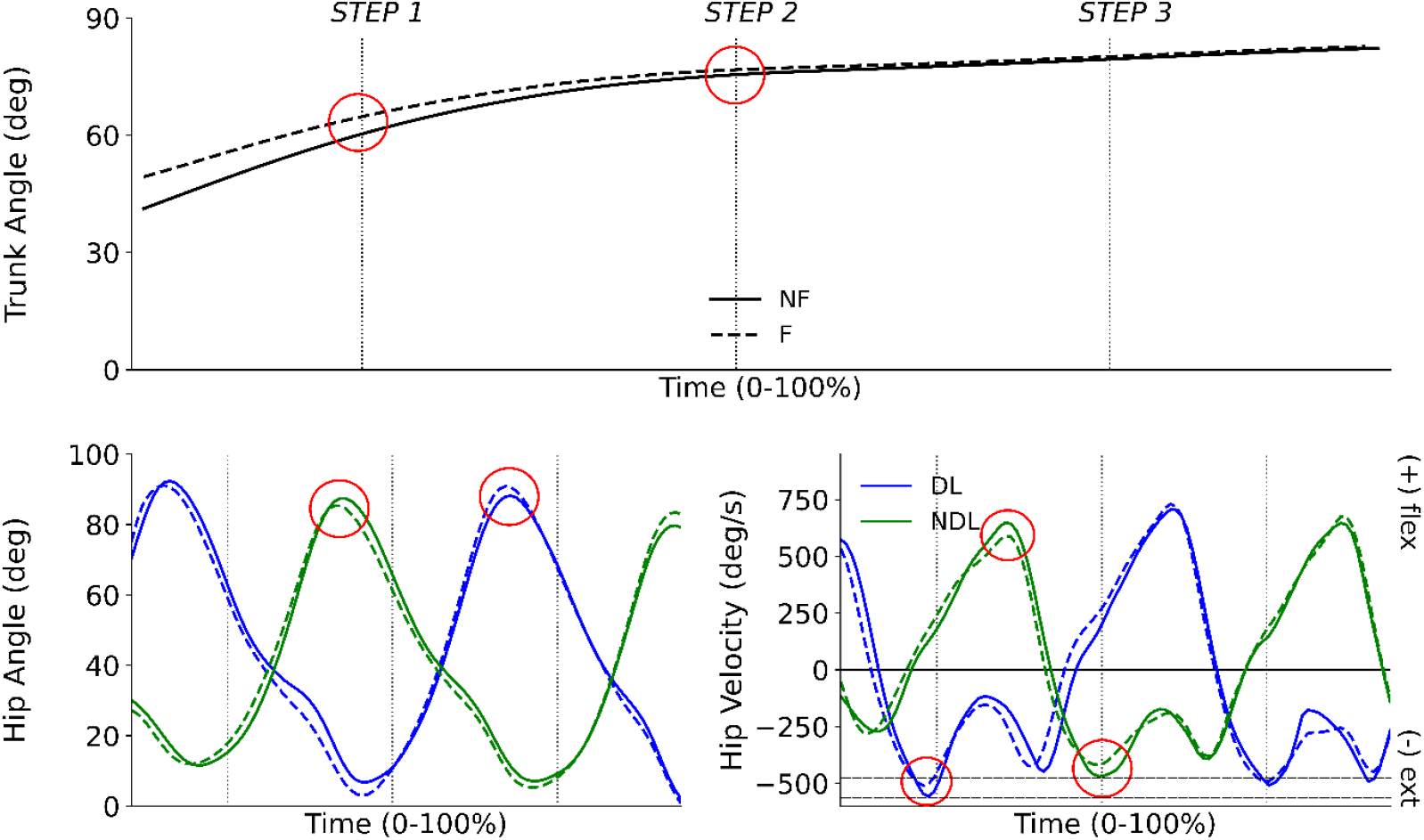
Trunk and hip kinematics during non-fatigued and fatigued acceleration trials. After fatiguing exercise, larger trunk angle relative to the ground were observed, i.e., a more upright posture during S1 and S2 (top); simultaneous decreases were detected in peak hip flexion angle and hip extension velocity for S2, while S1 and S3 was maintained (bottom right). Vertical dotted lines represent foot-strikes for steps 1, 2, and 3, respectively. Top: sagittal plane angle of the trunk relative to the ground (degrees) in non-fatigued (solid line) and fatigued (dashed line) conditions; bottom left: sagittal hip angle (degrees) for dominant leg (DL, blue) and non-dominant leg (NDL, green) in non-fatigued (solid line) and fatigued (dashed line) conditions from early protraction to toe-off for all three steps; middle right: sagittal hip velocity (deg/s) for DL (blue) and NDL (green) in non-fatigued (solid line) and fatigued (dashed line) conditions from early protraction to toe-off for all three steps. The time-dependent paired t-values of the SPM were set at p < 0.05, red circles indicate regions with statistical difference between non-fatigued and fatigued trials.

**Figure 4.**
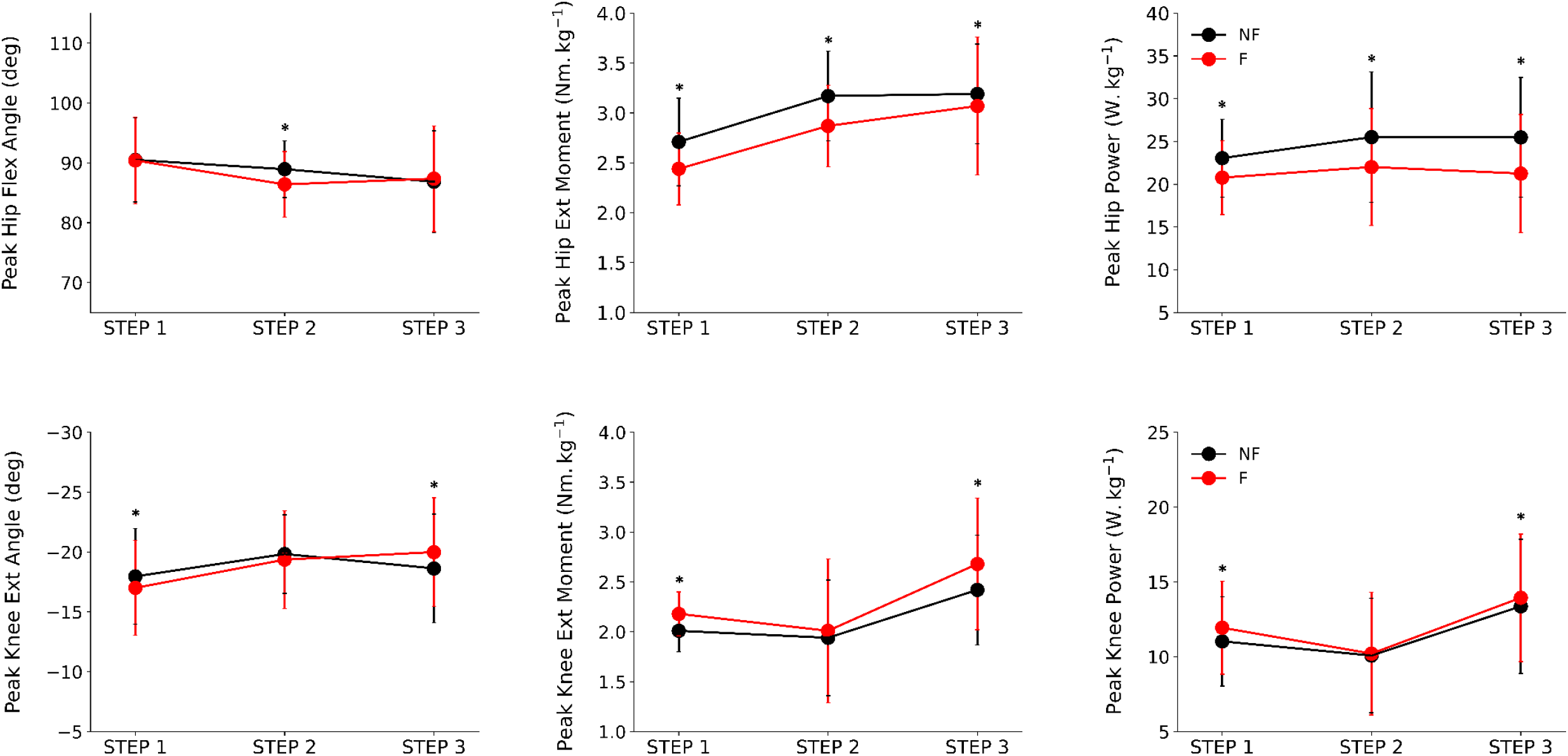
Peak hip and knee joint angles, moments, and positive power during non-fatigued and fatigued acceleration trials. After fatiguing exercise, peak hip extension moments and positive power decreased across all steps; peak knee extension moments and positive power increased for S1 and S3. Top left: peak hip flexion angle (degrees); top middle: peak hip extension moment (Nm.kg^-1^); top right: peak positive hip joint power (W.kg^-1^); bottom left: peak knee extension angle (degrees); bottom middle: peak knee extension moment (Nm.kg^-1^); bottom right: peak positive knee power (W.kg^-1^) during non-fatigued (black) and fatigued (red) conditions. *Statistical difference (*t*-test) between non-fatigued and fatigued trials (p < 0.05).

Peak hip extension moment and power significantly increased from S1 to S2, but no changes were detected between S2 and S3 (see Figure 3). Peak knee extension moment and power were similar from S1 to S2 but significantly increased from S2 to S3. Small increases in peak plantarflexion moment and power were observed with each successive step but did not reach statistical significance. Positive work was greatest at the ankle joint for S3 (38.6%), while similar contributions were made by the hip (S1: 33.9%; S2: 37.3%) and ankle (S1: 35.1%; S2: 36%) joints in S1 and S2. The hip work contribution in S3 (35.3%) was the next greatest, while the knee contributed least to all steps, with this difference being most marked in, and not different between, S2 and S3 (S1: 30.9%; S2: 26.7%; S3: 26.1%). Total positive work performed was similar for S1 (2.78 J·kg^-1^) and S2 (2.62 J·kg^-1^) but increased by 48% from S2 to S3 (to 5.10 J·kg^-1^).

### Step-to-step variable comparison between non-fatigued and fatigued accelerated running

Fatiguing exercise did not detectably alter either CoM horizontal velocity or vertical CoM displacement in any step. However, while ground contact times did not significantly change for S1, ∼6% increases were detected in S2 and S3 (with no significant difference between S2 and S3 – see Table 1). During S1, the horizontal position of the foot relative to the CoM at ground contact was unchanged by fatigue, but the foot landed further in front of the CoM in S2 (20%) and S3 (23%) compared to the non-fatigued condition. Horizontal foot velocity was also unchanged in S1 but tended to increase in S2 (26%) and S3 (15%) with fatigue, although statistical significance was not reached (p = 0.07 and p = 0.06, respectively). Vertical foot velocity and net horizontal impulse remained unchanged, but vertical impulse significantly increased ∼4-6% after fatigue across all steps (Figure 2).

### Kinematic, kinetic, and mechanical work comparison between non-fatigued and fatigued accelerated running

Sagittal trunk angle relative to the horizontal significantly increased ∼15% at foot ground contact during S1 and ∼8% in S2 with fatigue but was not different in S3 (Figure 3). The peak hip flexion angle also remained unchanged in S1 and S3 but decreased ∼3% for S2 (p = 0.03). Peak knee extension angle prior to foot-ground contact, i.e., the phase between early protraction and foot-ground contact, tended to increase for both S1 (5%) and S2 (2%) but decreased for S3 (7%); however, these mean changes did not reach statistical significance for any steps (Figure 3). Additionally, kinematics about the ankle joint did not change statistically in any step after fatigue.

Peak hip extension moments (S1: 12%, S2: 9%, S3: 4%) and power (S1: 10%, S2: 12%, S3: 17%) significantly decreased with fatigue, while knee moments and power significantly increased in S1 (∼7%) and S3 (∼4%) without change in S2, and peak plantarflexion moment increased only in S1 (8%). The distribution of positive work performed did not significantly change during S1 and S3, however the hip extensor contribution significantly decreased (∼4%) while knee extension contribution increased (∼4%) during S2. The total positive work performed did not change with fatigue, however significantly more work was performed in S1 (2.82 J·kg^-1^) than S2 (2.62 J·kg^-1^) and more positive work was performed in S3 (5.21 J·kg^-1^) than S2.

### Single-leg vertical jumps

Paired t-tests identified significant differences in jump height (*p* < 0.001) between DL and NDL in the non-fatigued condition, with DL ∼1.6 cm greater jump height and ∼8% greater vertical net impulse than NDL. After fatiguing exercise, no statistical differences were observed between DL and NDL (*p* < 0.069; *p* < 0.111). Jump height significantly decreased between non-fatigued and fatigued conditions for DL (*p* < 0.001) and NDL (*p* < 0.001)

## Discussion

### Overview

Contrary to the tested hypotheses, step-to-step velocity (i.e., rate of acceleration) did not decrease within the first three steps after the completion of fatiguing running exercise despite significant decreases in one-leg jump height being induced by the exercise and this same exercise previously leading to significant reductions in maximum running speed in the same cohort of athletes (Vial, Cochrane Wilkie, Turner & Blazevich, 2021). After performing fatiguing exercise, a more upright posture was adopted, slightly longer step lengths were used with the foot contacting the ground further in front of the CoM, longer ground contact times evolved, and greater vertical impulse was produced whilst horizontal impulse was maintained. In combination with differences in posture and step characteristics, hip flexion and extension velocity, and hip extension power and moments decreased but peak knee moments, power, and positive joint work increased, whilst ankle joint kinetics did not detectibly change with fatigue. Although these acute adaptations may speculatively help to maintain step-to-step velocity, it may also indicate that the hip extensors (i.e., gluteal and hamstring muscle groups) were more susceptible to fatigue than those surrounding the ankle and knee joints, and therefore contributed to a lesser extent during fatigued accelerated sprinting. Therefore, a strategy appears to have been adopted that incorporated longer ground contact times, a more upright posture, and greater peak knee moments and positive work production, which was sufficient to maintain the rate of acceleration during the first steps of acceleration after running-induced fatigue as before the fatiguing exercise.

The ability to retain acceleration rate observed after fatiguing exercise might have been gained through our cohort of athletes regularly performing repeated acceleration efforts, particularly under fatigued conditions, in both training and match situations. Team-sport athletes rarely run uninterrupted for long distances and seldom reach maximal sprinting speed, and therefore accelerated sprints are one of the most frequently completed high-speed running tasks in training and matches (Andrzejewski et al., 2018). This experience might speculatively have resulted in them developing, i.e., learning, to alter their movement patterns so as to maintain acceleration in the face of fatigue. This hypothesis requires more explicit scrutiny in future studies.

### Non-fatigued accelerated sprint performance and technique

During non-fatigued sprint acceleration, ground forces were primarily applied backwards and downwards to overcome the body’s inertia, and horizontal impulse was greatest during the first step but decreased with each successive step. Because inertial resistance decreases and momentum (mass × velocity) increases (Cavanagh, 1964), net force (horizontal direction) should decrease with the increase in velocity. This effect was observed in the present study, with ground contact times and horizontal foot velocity (i.e., forward foot velocity relative to CoM) decreasing as posture became more upright and resulted in the first step having the greatest net horizontal impulse, which decreased with each successive step. Although whole-body velocity increased at each step, vertical impulse was relatively unchanged across steps, likely due to inertial resistance decreasing whilst the gravitational force acting on the body remained constant across steps. Thus, the same vertical force was produced but with progressively shorter ground contact times. Although horizontal force decreased with each step, overall acceleration mechanics for steps 1 (dominant leg; DL), 2 (non-dominant leg; NDL), and 3 (DL again) remained similar. Equal relative positive work was performed at the hip and ankle joints in S1 and S2, while a slightly greater relative ankle joint contribution was observed in S3 (Figure 5). As such, we speculate that there was relatively equal contribution from the larger proximal hip extensors (i.e., gluteal and hamstring muscles) and distal elastic powered ankle joint (i.e., Achilles’ tendon, gastrocnemii, and soleus muscles) for propulsion during all steps. In agreement with previous work, maximum sprinting acceleration in non-fatigued conditions appears to largely depend on the work performed by the hip and ankle joints (Schache et al., 2019).

**Figure 5.**
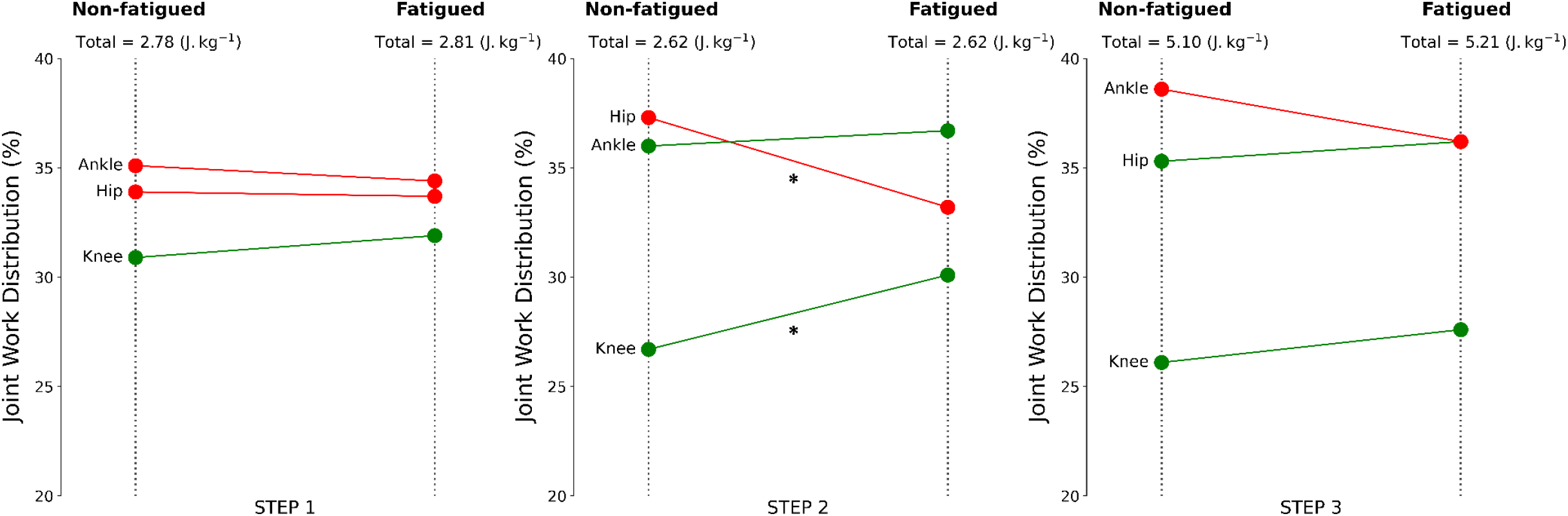
Joint work distribution of lower limb joints expressed as a percentage of total positive work across each step during non-fatigued and fatigued acceleration trials. For S2, positive hip joint work performed decreased while positive knee joint work increased (centre); no significant differences were observed for S1 (left) and S3 (right). Green line represents an increase from non-fatigued to fatigued conditions, whilst red line indicates a decrease. Total positive joint work (J kg^-1^) performed for each step is provided above each condition. *Statistical difference (*t*-test) between non-fatigued and fatigued trials (p < 0.05).

The individual limb contribution to total positive joint work was slightly greater in S1 than S2; however, a 48% increase was observed from S2 to S3. That is, the positive joint work performed in S3 was almost double that of S2, but horizontal velocity continued to increase. Intuitively, the positive joint work performed should reflect increases in whole-body velocity. For example, the relative joint work done may be expected to increase simultaneously with horizontal velocity. Yet, step-to-step velocity increased despite a lack of increase in total positive joint work done in S2. One explanation might be provided by the differences in posture between steps; a more forward leaning posture was adopted in S1 and S2 than S3, allowing the gravitational force to do work on the body to assist horizontal acceleration. The more forward position of the body’s centre of mass over the support (ground) leg results in more forward, rotational acceleration of the body over the foot, and thus forward acceleration of the body. Nevertheless, horizontal CoM velocity reached ∼1.51 m/s for S1, increased by 1.06 m/s to 2.57 m/s for S2, and increased 1.17 m/s to 3.74 m/s for S3; hence, the rate of acceleration was lowest in S2 (see Figure 1). The main findings of interest during non-fatigued accelerated sprinting included a non-linear decrease in acceleration with each step, with the third step producing nearly double the positive joint work of the previous steps. The change in step-to-step horizontal velocity was greatest for step 1 (1.51 m/s), decreased ∼30% to step 2 (1.06 m/s), then increased slightly to step 3 (1.17 m/s). These data, in combination with positive joint work done and jump test results (i.e., SLVJ jump height), indicate that the non-dominant leg was indeed the weaker limb.

### Non-fatigued and fatigued accelerated sprint performance and technique comparison

Unlike top speed sprinting, which requires large vertical forces to be produced in a very short foot-ground contact time of ∼0.1 s (Clark et al., 2017), accelerated sprinting requires a magnitude of vertical impulse that is sufficient to project the body into the next step with adequate time to reposition the leg towards the front of the body (Hunter et al., 2005). We observed that ground contact times during S1 were unaltered after fatiguing exercise compared to non-fatigued running but were increased in S2 and S3. Meanwhile, net horizontal impulse remained unchanged, and instead, significant increases in vertical impulse were observed across all steps. In addition, hip flexion and extension velocities decreased in the fatigued running condition with simultaneous decreases in hip extension moments and powers across all steps (i.e., dominant and non-dominant legs). Hunter et al. (2005) hypothesised that vertical impulse may be smaller during initial acceleration in individuals who are able to reposition their limbs more quickly in preparation for the next step. However, we observed a significant increase in vertical impulse after fatiguing exercise and, according to Hunter et al’s (2005) premise, the capacity to reposition the lower limbs (i.e., reduced hip flexion/extension velocity) in preparation for the next step may have been diminished due to fatigue. With an increase in vertical impulse, one might expect a simultaneous increase in vertical CoM displacement, however we instead observed a significant increase in the projection angle (i.e., a more upright posture - see Figure 3). Given the above, we speculate that in order to overcome the slowing of hip flexion and extension, the use of a more upright posture allowed more time to reposition the lower limbs for the following step, and therefore orientated the body in a better position to generate vertical impulse. That is, producing downward force is far easier if the body is more upright, and therefore vertical force production increased because of the change in posture.

In combination with greater vertical force production, we also observed a shift in joint contribution, including a decrease in hip joint contribution and simultaneous increase in peak knee joint moments, power, and positive work done in particular, after fatiguing exercise. However, slightly different strategies were used for dominant and non-dominant limbs. Greater peak knee extension moments and power were observed in S1 and S3 (i.e., dominant leg), but similar positive work was performed in the non-fatigued condition. While peak knee joint moments and power were similar for S2 (i.e., non-dominant leg) to the non-fatigued condition, positive hip joint work decreased whilst positive knee work increased. The longer ground contact times and greater vertical force production might have resulted from increased peak knee extension moments (dominant leg) and positive knee joint work performed (non-dominant leg). In prolonged, middle-distance speed running (Willer et al., 2021), a proximal shift from the ankle to the knee joint was observed with fatigue. Instead, we observed a relative reduction in positive joint work at the hip and increase at the knee joint. While the present results and those of Willer et al (2021) tend to reveal similar outcomes (i.e., a shift to knee joint work production), the point of difference is the proximo-distal direction of the shift. The relative shift from the hip extensors, which contribute significantly to horizontal force production in sprinting acceleration, indicates that fatigue might manifest differently in slower, continuous running, than multi-directional higher intensity exercise. An alternative explanation is that fatigue simply manifests differently between sprinting and jogging. Incorporating the use of periods of acceleration and sprinting in our fatiguing protocol, which requires significant hip work and power production, might influence this capacity in a subsequent test of acceleration, whereas continual jogging may evoke greater distal muscle fatigue that is then detected during a subsequent jogging test. Irrespective of the mechanism or site of fatigue, our cohort of athletes adapted their movement pattern to negate acceleration loss.

While one hypothesis may be that our cohort have a learned ability to shift muscle force production from more fatigued to less fatigued muscles to maintain the rate of acceleration, an alternative explanation, or additional benefit, is that the hip-to-knee muscle force redistribution may serve a fatigued-induced muscle injury protective purpose, particularly for the injury prone hamstring muscles. Hamstring muscle injuries are highly prevalent in modern sports and are the most prevalent injury in running-based sports such as Association Football (soccer) (Ekstrand et al., 2011; Woods et al., 2004). Although injuries are problematic for modern athletes, injury in early humans would have had severe consequences, so the adoption of injury reduction practices or preserving acceleration ability to evade predators whilst fatigued, would have been important for survival. A shift from hip extensor (hamstring) to knee extensor (quadriceps) work and power production may provide a useful injury minimisation purpose, especially since hip flexion angles are acute, and thus places the hamstring muscles at long length, when the body has a significant forward lean during accelerative running. This hypothesis is worthy of explicit examination in future studies.

### Summary

In the non-fatigue trials, the first steps of accelerated sprinting appear to mostly depend upon hip and ankle work and power contributions. Our primary findings included a non-linear decrease in the rate of acceleration, with step 2 (i.e., non-dominant leg) having the lowest rate of acceleration. Secondly, similar positive work was done during the first (DL) and second (NDL) steps, while work in the third step (DL) was nearly double that of S1 and S2. These data, in combination with jump test results (i.e., SLVJ jump height), indicate that the non-dominant leg is indeed the weaker leg. With fatigue, we observed a more upright posture, longer ground contact times, increased vertical impulse, and a shift in peak moments, power, and positive work done from the hip to the knee joint. We theorise that this shift is either a direct cause or a result of reduced hip flexion and extension velocity to reposition the lower limbs in preparation for the next step. Similar to previous research in slower, more economical running, we observed that running-induced fatigue led to a compensatory increase in knee joint work (Willer et al., 2021). However, unlike those previous findings, fatigue promoted a shift from the larger, powerful hip extensors rather than elastic powered ankle propulsion to muscle-dominant knee power production. Thus, maximal acceleration, contrary to slower-speed running, does not seem to be compromised by a loss of ankle work or power whilst in a fatigued state. As an alternative, we suggest that fatigue induced through the combination of slow, moderate, and high speed multi-directional movements used in the present study may have induced a marked hip extensor fatigue (i.e., gluteal and hamstring muscle groups) with a compensatory increase in knee peak moments, power, and positive work (i.e., by quadriceps muscles); the degree of hip extensor fatigue evoked by the running protocol requires quantification in future experiments. Regardless, this adaptation may be used as a strategy to maintain the rate of acceleration or reduce the risk of injuring muscles (e.g. hamstrings) that may be more susceptible to fatigue. Nevertheless, the results provide insight into the mechanics that hunter-gather people may have also used when undertaking similar accelerating activities, particularly when fatigued. However, it is noteworthy that testing was conducted in an indoor laboratory on a synthetic athletics track while the fatiguing protocol was completed outdoors on a grassed sports field. Future research should investigate accelerative running mechanics on grassed and dirt surfaces, which may be more comparable to the surfaces used in hunter-gatherer societies.

## Author Contributions

SV, JCW, and AJB designed research; SV and MT performed research; SV and AJB analysed data; and SV, JCW, MT, and AJB wrote the paper.

